# Transfer of IgG from Long COVID patients induces symptomology in mice

**DOI:** 10.1101/2024.05.30.596590

**Authors:** Hung-Jen Chen, Brent Appelman, Hanneke Willemen, Amelie Bos, Judith Prado, Chiara. E. Geyer, Patrícia Silva Santos Ribeiro, Sabine Versteeg, Mads Larsen, Eline Schüchner, Marije K. Bomers, Ayesha H.A. Lavell, Amsterdam UMC COVID-19 biobank, Braeden Charlton, Rob Wüst, W. Joost Wiersinga, Michèle van Vugt, Gestur Vidarsson, Niels Eijkelkamp, Jeroen den Dunnen

**Affiliations:** Center for Experimental and Molecular Medicine, Amsterdam Institute for Infection and Immunity, Amsterdam University Medical Centers, location AMC, Amsterdam, Netherlands; Center for Translational Immunology, University Medical Center Utrecht, Utrecht University, Utrecht, Netherlands; Department of Experimental Immunohematology, Sanquin Research, Amsterdam, Netherlands; Faculty of Behavioral and Movement Sciences, Vrije Universiteit, Amsterdam, Netherlands; Department of Infectious Diseases, Amsterdam Institute for Infection and Immunity, Amsterdam University Medical Centers, location VUMC, Amsterdam, Netherlands

## Abstract

SARS-CoV-2 infections worldwide led to a surge in cases of Long COVID, a post-infectious syndrome. It has been hypothesized that autoantibodies play a crucial role in the development of Long COVID and other syndromes, such as fibromyalgia and myalgic encephalomyelitis/chronic fatigue syndrome (ME/CFS). In this study, we tested this hypothesis by passively transferring total IgG from Long COVID patients to mice. Using Glial Fibrillary Acidic Protein (GFAP) and type-I interferon expression, we stratified patients into three Long COVID subgroups, each with unique plasma proteome signatures. Remarkably, IgG transfer from the two subgroups, which are characterized by higher plasma levels of neuronal proteins and leukocyte activation markers, induced pronounced and persistent sensory hypersensitivity with distinct kinetics. Conversely, IgG transfer from the third subgroup, which are characterized by enriched skeletal and cardiac muscle proteome profiles, reduced locomotor activity in mice without affecting their motor coordination. These findings demonstrate that transfer of IgG from Long COVID patients to mice replicates disease symptoms, underscoring IgG’s causative role in Long COVID pathogenesis. This work proposes a murine model that mirrors Long COVID’s pathophysiological mechanisms, which may be used as a tool for screening and developing targeted therapeutics.

## Introduction

The emergence of Coronavirus Disease 2019 (COVID-19), caused by Severe Acute Respiratory Syndrome Coronavirus 2 (SARS-CoV-2), has sparked a global health crisis, with over 780 million cases and 7 million deaths reported to date [1]. Ample evidence highlights a concerning trend among COVID-19 survivors, with a significant subset (> 10%) experiencing a spectrum of persistent symptoms exceeding 12 weeks post initial recovery [2–4]. This condition is referred to as Post-Acute Sequelae of SARS-CoV-2 infection (PASC), post-COVID syndrome, or colloquially as Long COVID. Long COVID presents a heterogeneous array of symptoms, including cough, fatigue, post-exertional malaise (PEM), neurocognitive impairment (brain fog, sleep and anxiety disorders), sensory and musculoskeletal manifestations (joint pain, chest pain, muscle-ache), postural orthostatic tachycardia syndrome, fever, shortness of breath, gastrointestinal disturbances, and palpitations [3, 5–7]. However, the underlying pathophysiology of Long COVID remains elusive.

Several potential mechanisms have been proposed, connecting Long COVID symptomatology to dysregulated interferon (IFN) response, inflammation, cellular metabolism, persistent infection, dysbiosis, neuroinflammation, and autoimmune processes [3, 8–14]. Among these factors, studies have demonstrated that autoimmunity is induced in both acute and post-acute phases of COVID-19 [15–17]. While acute disease severity correlates with anti-viral protein antibodies [18], Long COVID is characterized by the presence of autoantibodies targeting diverse self-antigens [19]. These Long COVID-associated autoantibodies bind to chemokines [20], G protein-coupled receptors [21], neurotransmitters [22], and various immunomodulating proteins [16, 23]. Moreover, it has been hypothesized that autoantibodies play a crucial role in other chronic fatigue and post-acute infection syndromes, such as post-Lyme disease syndrome, Q-fever fatigue syndrome, fibromyalgia, and myalgic encephalomyelitis/chronic fatigue syndrome (ME/CFS) [6]. Yet, whether these autoantibodies are mere bystanders or active contributors to Long COVID symptoms is not known.

Recent experimental evidence supports the potential involvement of autoantibodies in driving Long COVID symptomatology. Following therapeutic apheresis, clinical improvement in Long COVID patients appears to be associated with autoantibody reduction [22]. Moreover, transfer of immunoglobulin G (IgG) from fibromyalgia patients to mice induces pain associated behavior [24]. Hence, we postulate that autoantibodies may play a causal role in the manifestation of Long COVID symptoms in at least a subset of patients. In this study, we set out to elucidate the involvement of autoantibodies in Long COVID pathogenesis by establishing a patient-derived IgG-transferring mouse model for Long COVID.

## Methods

### Study population

This study comprised Long COVID patients seen at the Amsterdam UMC outpatient Post COVID-19 Clinic. All patients were seen by a clinician, dedicated to post COVID-19, and had been diagnosed with Long COVID according to the WHO criteria (the continuation or development of new symptoms three months after SARS-CoV-2 infection, with these symptoms lasting for at least two months with no other explanation) and were required to have a reduction in working hours after SARS-CoV-2 infection. For the current study we selected patients between 18-65 years old who had previously a proven mild SARS-CoV-2 infection (non-hospitalized). Venous blood was obtained in the Amsterdam UMC post-COVID-19 biobank study, a minimum of 90 days after initial SARS-CoV-2 infection. The Amsterdam UMC post-COVID-19 Biobank study was approved by the institutional biobank ethics committee (Amsterdam UMC 2020_065). Demographics, comorbidities, symptomology and medications were derived from electronic health records. For subgroup analysis three Long COVID groups were created based on <u>Meso Scale Discovery (</u>MSD) analysis. For this study we used two different control groups; healthy control subjects sampled prior to the SARS-CoV-2 pandemic and healthy control subjects’ samples after mild SARS-CoV-2 infection but without residual symptoms (S3 study, NL73478.029.20, n=15) [25]. All blood samples were processed, aliquoted and frozen within 4h of blood draw. Written informed consent was obtained for all study participants.

### Meso Scale Discovery (MSD) multiplex assay

U-PLEX and R-PLEX Custom Human Cytokine assays were employed for the detection of IL-1β, IL-6, IL-10, TNF, IFN-α2a, IFN-β, IFN-γ, GFAP, neurofilament L (NFL), and total Tau. The analysis was performed on EDTA plasma (EDTA tube, 2000g for 10 min) of 34 Long COVID and 15 healthy controls. The lyophilized single or cocktail mix calibrators were reconstituted in provided assay diluents. MSD plates were prepared by coating them with supplied linkers and biotinylated capture antibodies as per the manufacturer’s instructions. The assays were conducted according to the manufacturer’s protocol, with the undiluted plasma samples and standards incubated overnight at 4°C. Electrochemiluminescence signals were measured using a MESO QuickPlex SQ 120 plate reader (MSD) and analyzed using Discovery Workbench Software (v4.0, MSD). The concentration of each sample was determined using a four-parameter logistic model generated with the standards, and the concentrations were calculated based on the certificate of analysis provided by MSD. Concentrations below the lower limit of detection were imputed as half of the lowest detected value.

### OLINK proteomics

Olink Proteomics technology was employed for protein profiling analysis. Due to limited samples available, the analysis was performed on EDTA plasma (EDTA tube, 2000g for 10 min) of 31 out of 34 Long COVID patients. Per protocol, samples were randomized across plates and run alongside a negative control (buffer), plate control, and a sample control. A total of 2,944 proteins were measured, with 2,865 proteins quantified after quality check filtering. For data normalization we employed the normalizeVSN and normalizeQuantiles packages. Normalized protein expressions (NPX) are reported as log2 values as per OLINK protocol.

### Human IgG purification

IgG was purified from 800 μl of patient serum or healthy donor EDTA plasma using Protein G-conjugated beads (Cytiva cat# 17061801). The beads were added to 1-ml gravity flow columns (Thermo Scientific, 89896) and washed with PBS. Serum/plasma was diluted 1:1 with PBS (pH = 7) and added to the washed column. The flow through was collected and reapplied to the column five times to increase recovery rate. After washing, bound IgG was eluted with elution buffer (0.1 M glycine buffer pH = 2.7) and immediately neutralized with neutralization buffer (1 M Tris buffer pH = 9). Protein concentration was determined by nanodrop measurement. Salt content was removed with Slide-A-Lyzer MINI Dialysis Device (Thermo Scientific). Purity of the eluted fractions was determined via SDS page and WB. Prior to injection, the samples were concentrated in saline buffer using Vivaspin to achieve a final concentration of 13 mg/ml.

### Mouse model and behavioral assays

Experiments were conducted using adult male and female (aged 8-16 weeks) C57BL/6 mice (Janvier laboratories). Mice were maintained in the animal facility of the University of Utrecht and housed in groups under a 12h:12h light-dark cycle, with food and water available *ad libitum*. The cages contained environmental enrichment, including tissue papers and shelter. All experiments were performed in accordance with international guidelines and approved by the local experimental animal welfare body and the national Central Authority for Scientific Procedures on Animals (CCD, AVD11500202010805). All mice were acclimatized for the behavioral assays before measurements. Baseline measurements were performed before the mice were given any injection. After the baseline measurements, mice were injected intraperitoneal with ∼6.5 mg IgG/mouse (260mg/kg), approximately 1/3 of the total circulating mouse IgG [26]. The following behavioral tests were performed:

Heat withdrawal latency times were determined using the Hargreaves test (IITC Life Science) [27]. Mechanical thresholds were determined using the von Frey test (Stoelting) with the up-and-down method previously described [28]. To minimize bias, animals were randomly assigned to the different groups prior to the start of experiment using Randomice software (v1.1.5, GitHub) based on the following variables: age, weight, cage mechanical sensitivity (Von Frey test) and locomotor activity (rotarod and open field tests) at baseline [29]. All experiments were performed by operators blinded to the treatments.

Local motor activity and stamina was determined with an open field test and rotarod analysis, respectively. Mice from different groups were tested interspersed throughout the trials. To measure local motor activity, one mouse at a time was placed in the center of an arena (30 cm × 15 cm) and spontaneous behavior was recorded for 30 min (video camera imagingsource DMK22AUC03) and analyzed (e.g. distance, immobility) using the ANY-maze software. To assess neurological deficits, like motor performance and stamina, mice were placed in the rotarod at a fix rotation of with accelerated speed [30]. Time to fall was recorded for 300s with 1) a fixed rotation of 12 rpm or 2) during the acceleration test which started at 4 rpm and increased overtime to 40 rpm. The open field arena and rotarod were cleaned thoroughly with a 5% alcohol/water solution between each mouse to minimize odor cues.

### Immunohistochemistry

Murine lumbar spinal cord and lumbar dorsal (L3-L5) root ganglia (DRG) tissue were embedded and frozen in optimal cutting temperature (OCT) freezing matrix (Sakura) using dry ice. Tissue slides of 10 µm thick were cut (Leica CM3050 S Cryostat) and kept at −80°C. On the day of staining, tissue slides were thawed at room temperature for 30 minutes and tissue was encircled using a Dako-pen. Slides were washed 3-times with 200 µl PBS, and fixated with 4% paraformaldehyde for 10 minutes. After washing slides with PBS with 0.3% Triton X-100 twice for 5 minutes. Tissue was blocked with 2% IgG-free BSA blocking buffer (ImmunoResearch cat:001-000-161). After tapping the blocking buffer from slides, they were incubated for 2 hours with 200 µl 1:250 goat anti-human IgG-Alexa fluor-647 in 3-time diluted blocking buffer at room temperature (ImmunoResearch Jackson cat:109-606-088). Subsequently, tissue slides were washed 3-times with diluted blocking buffer and further blocked with 200 µl 1:500 human IgG (nanogram, Sanquin) for 30 minutes. Slides were washed 1-time with diluted blocking buffer. Spinal cord sections were incubated with, mouse anti-GFAP (OriGene, BM2287; 1:200) overnight at 4°. The next day, slides were washed with 1:3 diluted blocking buffer 3-times for 1 minute. Subsequently, slides were incubated with anti-mouse Alexa fluor-594 antibody (Invitrogen, A21203; 1:500) for 90 minutes at room temperature. DRG sections were incubated with rabbit anti-glutamine synthesis (Abcam, ab73593; 1:200), mouse anti-GFAP (OriGene, BM2287; 1:200) and rat anti-F4/80 (Cedarlane, CL8940AP; 1:500) overnight at 4°C. The next day, slides were washed with 1:3 diluted blocking buffer (3-times, 1 minute) and incubated with anti-mouse Alexa fluor-594 antibody (Invitrogen, A21203), anti-rabbit Alexa fluor-750 (Invitrogen, A21039), and anti-rat Alexa fluor-488 (Invitrogen, A21208) all at 1:500 for 90 minutes at room temperature. After washing 3-times with PBS, slides were incubated with NeuroTrace™ 435/455 Blue Fluorescent Nissl Stain (Thermo Fisher, N21479; 1:300) for both spinal cord and DRG for 20 minutes. After washing sections were imbedded using FluoSafe (Merck, 345789) and kept at 4°C until imaging. Imaging was performed using a Thunder Wide Field Fluorescence microscope (Leica). The intensity and area of human IgG staining in mice was quantified using Image J software. The positive area for IgG was defined as the area with set pixel intensity and normalized to the total area of the section.

### Statistical analysis

For subject demographics and targeted biomarker analysis, histograms and Shapiro-Wilk tests were employed to assess data distributions and normality. Categorical values were depicted in absolute numbers alongside percentages in brackets. Parametric quantitative variables were shown as means ± standard deviation, while nonparametric quantitative variables were presented as median and interquartile ranges (25^th^ and 75^th^ percentiles). Categorical data were analyzed using Fisher’s exact test. Non-normally distributed data underwent Box-Cox transformation. Continuous parametric data were assessed using either a t-test analysis of variance, with Tukey HSD post-hoc testing applied when appropriate. Continuous nonparametric data were analyzed using the Mann-Whitney U test, Kruskal-Wallis H test, or pairwise Kruskal-Wallis test with Benjamini-Hochberg (BH) correction where appropriate. For Olink proteomics, partial least squares discriminant analysis (PLS-DA) and differential analysis for variance-stabilized quantile-normalized data were conducted using R packages mixOmics (v.6.24.0) and limma (v3.56.2). Principle components derived from PLS-DA were subjected to Gene Set Enrichment Analysis using R package fgsea (v.1.26.0). Protein-protein interaction analysis for differentially regulated proteins was carried out using the Metascape platform on 2024-02-06. In the behavioral assessment of mice, measured data was adjusted for pre-human IgG injection baseline measurements. Post-hoc comparisons of continuous data were performed using Empirical Mean Differences with BH adjustment with R package emmeans (v.1.10.1). A significance threshold of p < 0.05 was applied. The analyses and visualization were executed under R environment (v.4.3.2) or using GraphPad Prism (v.9.0).

## Results

### Long COVID patients are characterized by altered plasma levels of interferons and GFAP

A total of 34 patients were recruited from the Amsterdam UMC outpatient post-COVID-19 clinic. All participants had confirmed prior SARS-CoV-2 infection and were in good physical and mental health prior to infection. None of the individuals were hospitalized for COVID-19, and their symptoms persisted for a minimum of six months following the initial infection (Table 1). The diagnosis of Long COVID was made by a physician dedicated to post-COVID-19 at the Amsterdam UMC. Patients presented a varied array of symptoms, with fatigue being consistently reported among all participants. Additionally, 29 out of 34 patients experienced PEM, 25 out of 34 reported pain symptoms, and 26 out of 34 were unable to resume their previous occupational roles at the time of inclusion.

**Table 1.**
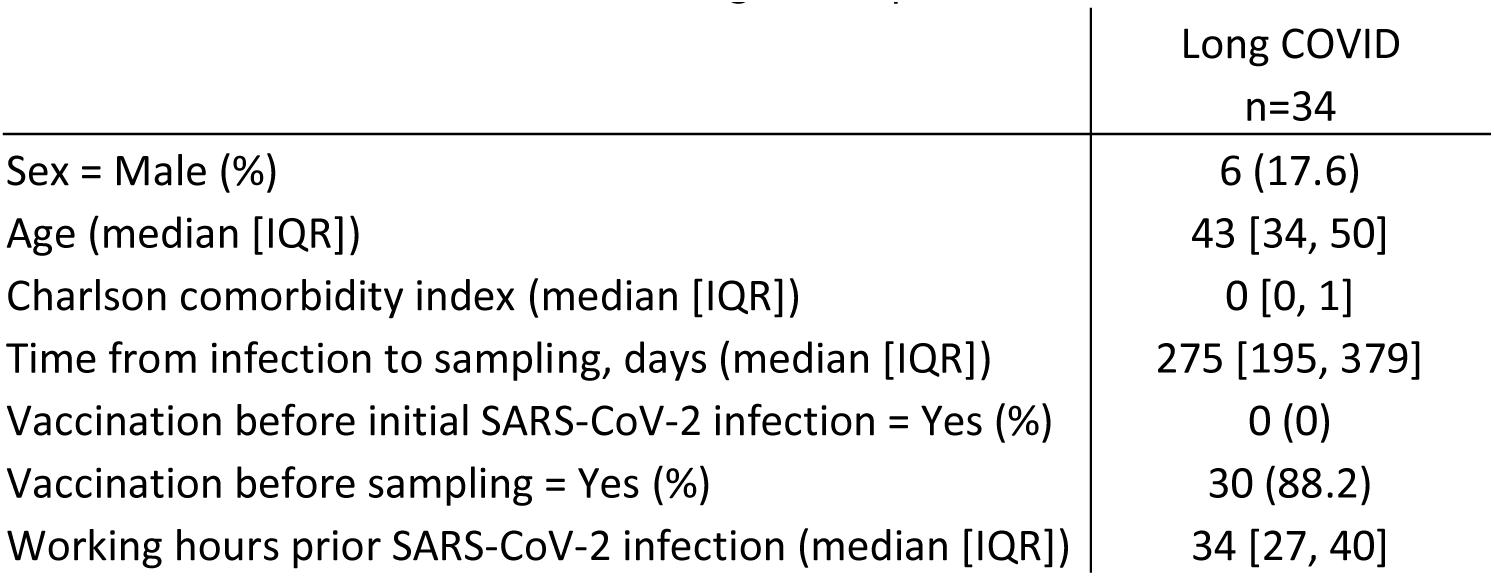
Baseline characteristics of Long COVID patients.

Since previous studies have shown that Long COVID is characterized by altered IFN levels [8, 31], chronic inflammation [32, 33], and signs of neuronal damage and neuroinflammation [34–36], we performed targeted biomarker quantification, measuring prototypic COVID-19 pro-inflammatory cytokines [37], type-I and -II IFNs, and neuronal damage and astrogliosis markers. In addition to the 34 Long COVID patients, we included age-, sex-, and post-infection time-matched 15 healthy controls (HC) who did recover from a prior mild SARS-CoV-2 infection without residual symptoms (Table S1). Plasma levels of acute-phase COVID-19 pro-inflammatory cytokines (IL-1β, IL-6, TNF, GM-CSF) were comparable between Long COVID patients and healthy controls (Supp. Fig. 1A-D). Interestingly, IFN-γ, the type-II IFN produced mainly by T cells and NK cells, was lower in Long COVID patients as compared to healthy controls (Fig 1A). Conversely, in Long COVID patients, IFN-β, a type-I IFN produced by most nucleated cells upon viral infection, tended to be elevated (Fig. 1B). The other type-I IFN, IFN-α2a (Fig. 1C), predominantly produced by plasmacytoid dendritic cells was not affected. Moreover, Glial Fibrillary Acidic Protein (GFAP), an astroglial activation marker, was detectable in 10 out of 34 Long COVID patients and significantly elevated in the patient group, whilst undetectable in all healthy controls (Fig. 1D). We did not observe significant differences in the neurodegenerative markers TAU and neurofilament light chain (NFL) between the patients and controls (Fig. 1E-F).

**Figure 1.**
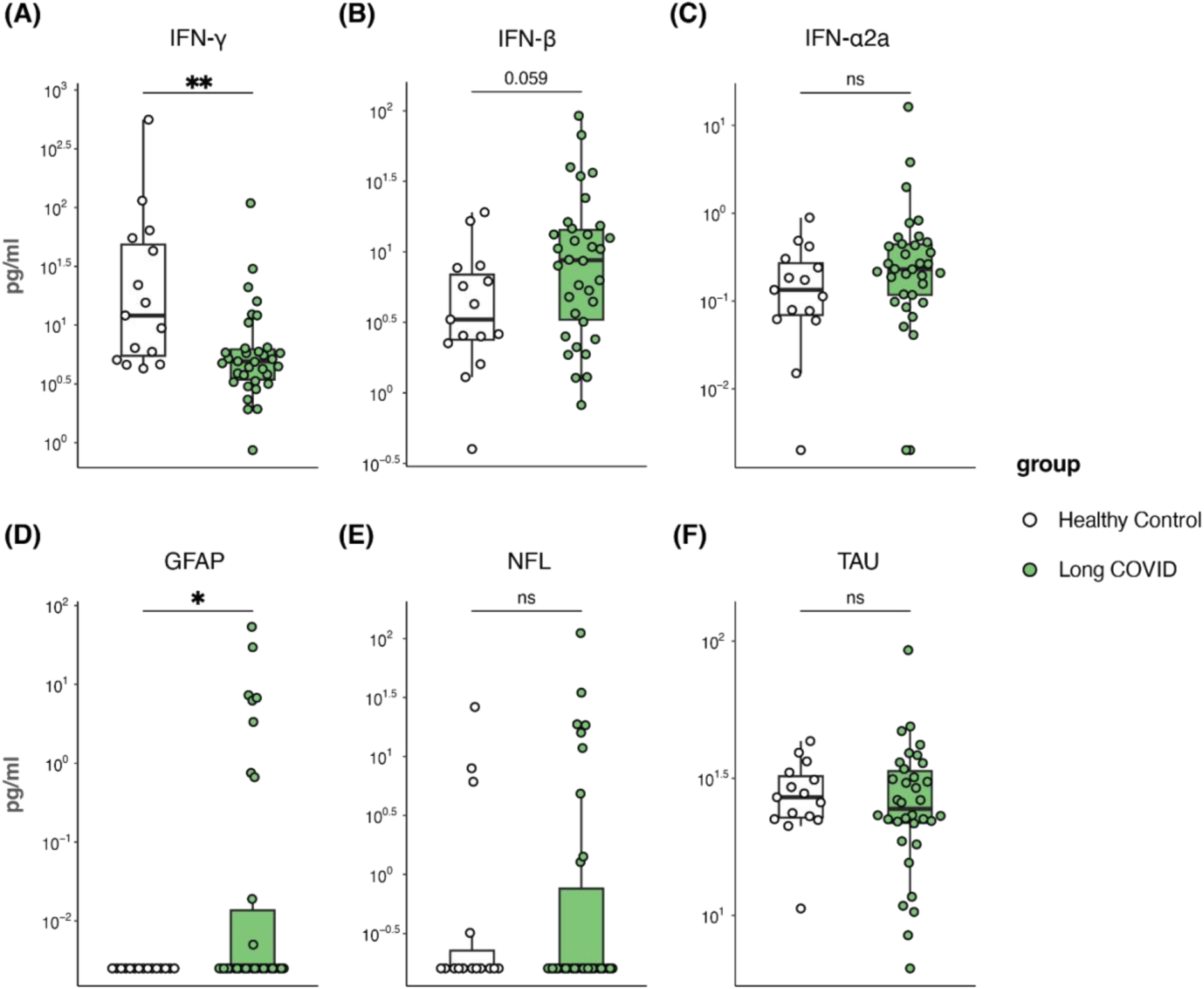
Plasma biomarkers of Long COVID compared to healthy SARS-CoV-2 recovered participants. Targeted quantitative measurements of plasma biomarkers of 34 Long COVID patients and 15 healthy controls with prior SARS-CoV-2 infection using MSD: **(A,B)** type-I interferons, **(C)** type-II interferon **(D)** astroglial activation marker Glial Fibrillary Acidic Protein (GFAP), neurodegenerative markers **(E)** neurofilament light chain (NFL) and **(F)** TAU. All values are shown in pg/ml on a logarithmic y-axis. *p<0.05; **p<0.01; ns, non-significant.

Next, we stratified Long COVID patients based on GFAP and IFN seral levels, given their disturbed concentrations in Long COVID patients and the known pathogenic roles of astrogliosis and type-I IFNs. We established three subgroups of patients. First, Long COVID-1 (LC-1) comprising 12 patients, characterized by elevated levels of neuronal damage and astroglia activation markers NFL, TAU, and GFAP (Fig. 2A-C). Subsequently, we divided the remaining patients into LC-2 and LC-3 based on type-I IFNs (Fig. 2D-E). LC-2 consist of 10 patients that had higher levels of IFN-α2a and IFN-β compared to group LC-3. Long COVID-3 (LC-3, 12 patients) have lower levels of TAU, type-I IFNs, and acute-phase pro-inflammatory cytokines IL-1β and IL-6 compared to LC-2 (Fig. S2A-D). Notably, we did not observed differences in IFN-γ levels between subgroups (Fig. 2F).

**Figure 2.**
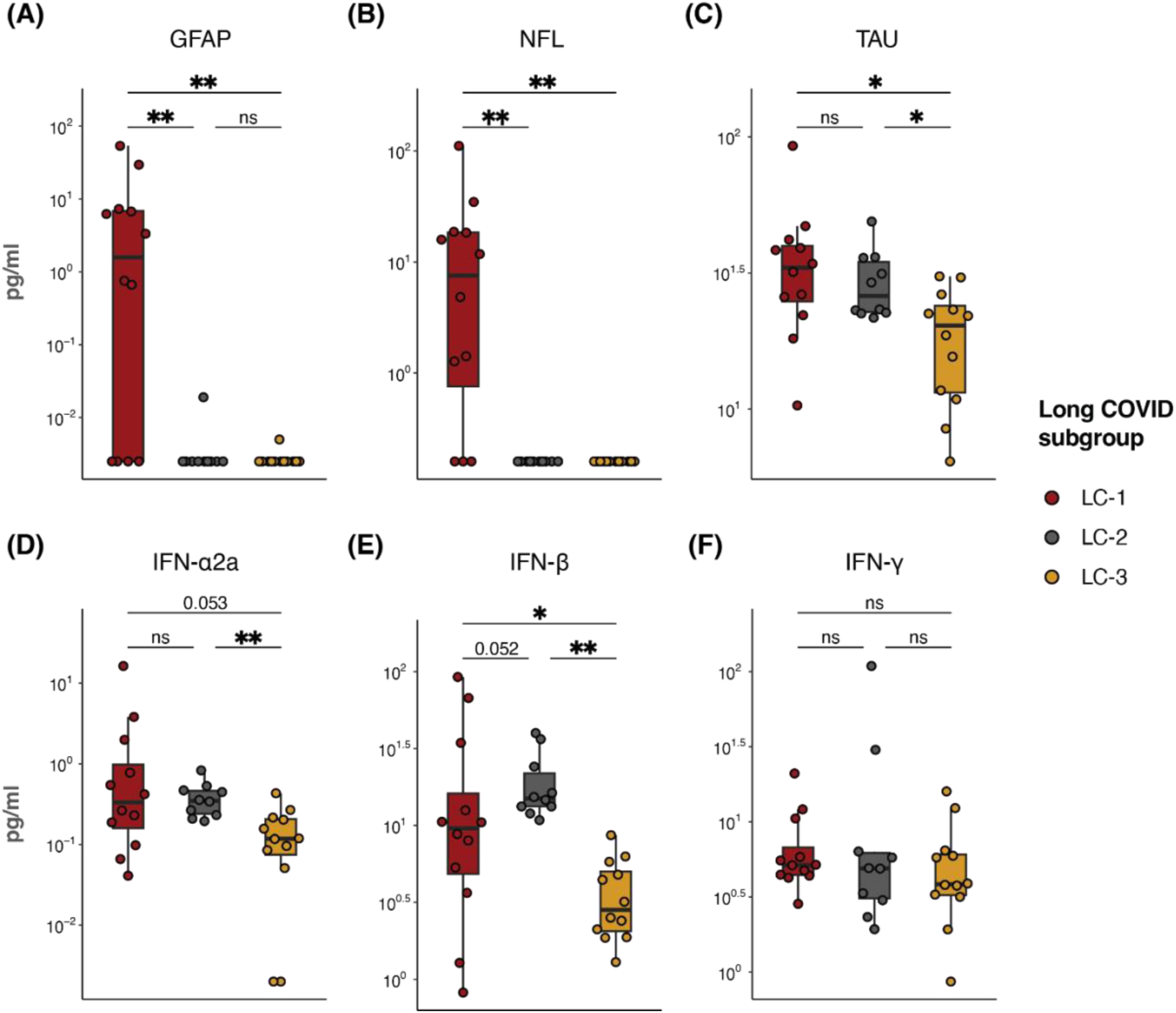
Plasma biomarkers in the Long COVID subgroups. Targeted quantitative measurements of astroglial activation marker Glial Fibrillary Acidic Protein (GFAP), neurodegenerative markers neurofilament light chain (NFL) and TAU, type-I and type-II interferons from 3 Long COVID (LC) patient subgroups (n = 12 for LC1, 10 for LC-2, 12 for LC-3). All values are shown in pg/ml on a logarithmic y-axis. *p<0.05; **p<0.01; ns, non-significant.

### Proteomics analysis reveals distinct pathways across Long COVID subgroups

Next, to explore potential shared pathogenesis within each subgroup, we conducted an extensive proteomics analysis, relatively quantifying the expression of 2865 proteins. We analyzed samples from 31 out of 34 long COVID patients, with 3 samples excluded due to limited sample volumes. We first employed supervised clustering with the Partial Least-Squares Discriminant Analysis (PLS-DA) algorithm to identify plasma proteins distinguishing the three subgroups. PLS-DA revealed a separation of LC-1 from LC-2 and LC-3 on principal component 1 (PC1), while LC-2 and LC-3 segregated on PC2 (Fig. 3A). We then performed Gene Set Enrichment Analysis (GSEA) on PC1 and PC2 to identify the pathways contributing to this separation. GSEA of PC1 indicated that LC-1 is associated with reduced plasma levels of cell surface proteins and elevated levels of intracellular transport proteins (Fig. 3B), whereas PC2 revealed specific enrichment of muscle-related proteins in LC-2 and higher levels of lipoproteins in LC-3 (Fig. 3C). By clustering the top 200 contributing components per PC in both PC1 and PC2, we identified several protein clusters differentially accumulated among the subgroups (Fig. 3D). For example, clusters C1, C2, and C3, containing neuronal damage and astroglia activation markers NFL and GFAP, were more abundant in LC-1 patients. Clusters C5 and C6, consisted of immune activation markers such as IL-12 (IL-12p35, IL-12p40) and HIF-1α that were higher in plasma of LC-2. Clusters C3, C4, and C6, containing immune cytokines and receptors like TNFSF9, IL4R, and TGFBR1, showed higher levels in LC-3.

**Figure 3.**
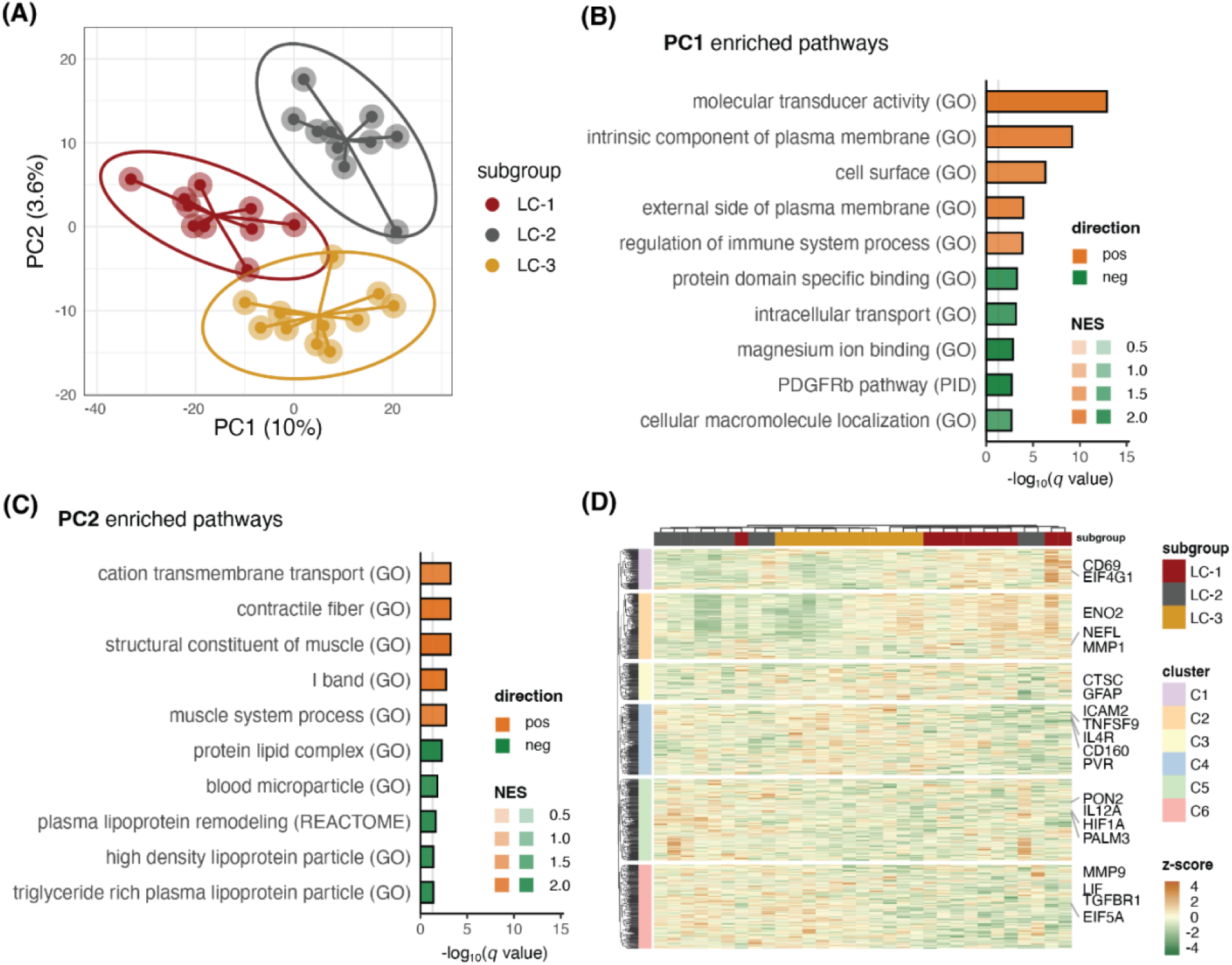
Plasma proteomics clustering analysis identified biomarkers distinguishing three Long COVID (LC) subgroups. Partial Least-Squares Discriminant Analysis revealed a separation of LC-1 from LC-2 and LC-3 on principal component 1 (PC1), while LC-2 and LC-3 segregated on PC2 **(A)**. Gene Set Enrichment Analysis results of PC1 **(B)** and PC2 **(C).** Clustering analysis of identified components in PC1-2 showed 6 protein clusters with distinct expression patterns **(D).**

To identify subgroup-specific alterations, we conducted differential analyses by comparing each subgroup with the other two. Consistent with our grouping strategy, we found increased levels of GFAP and NFL in LC-1 as well as elevated type-I IFN responding protein CXCL10 in LC-2 (Supp. Table 3). Next, we performed protein-protein interaction enrichment analysis for the differentially regulated proteins against the total proteins measured as enrichment background. In LC-1, interconnected protein modules were positively enriched in cytoskeleton, nervous system, and Golgi-related proteins, while there was a reduction in bacterial response and lysosome proteins (Fig. 4A). LC-2 exhibited higher muscle-related proteins and lower neurotransmitter proteins compared to LC-1 and LC-3 (Fig. 4B). Lastly, in LC-3, leukocyte activation proteins were elevated whilst muscle and neural proteins were reduced (Fig. 4C). These elevated muscle and neural proteins in plasma may indicate proteins released from local tissue damage entering circulation.

**Figure 4.**
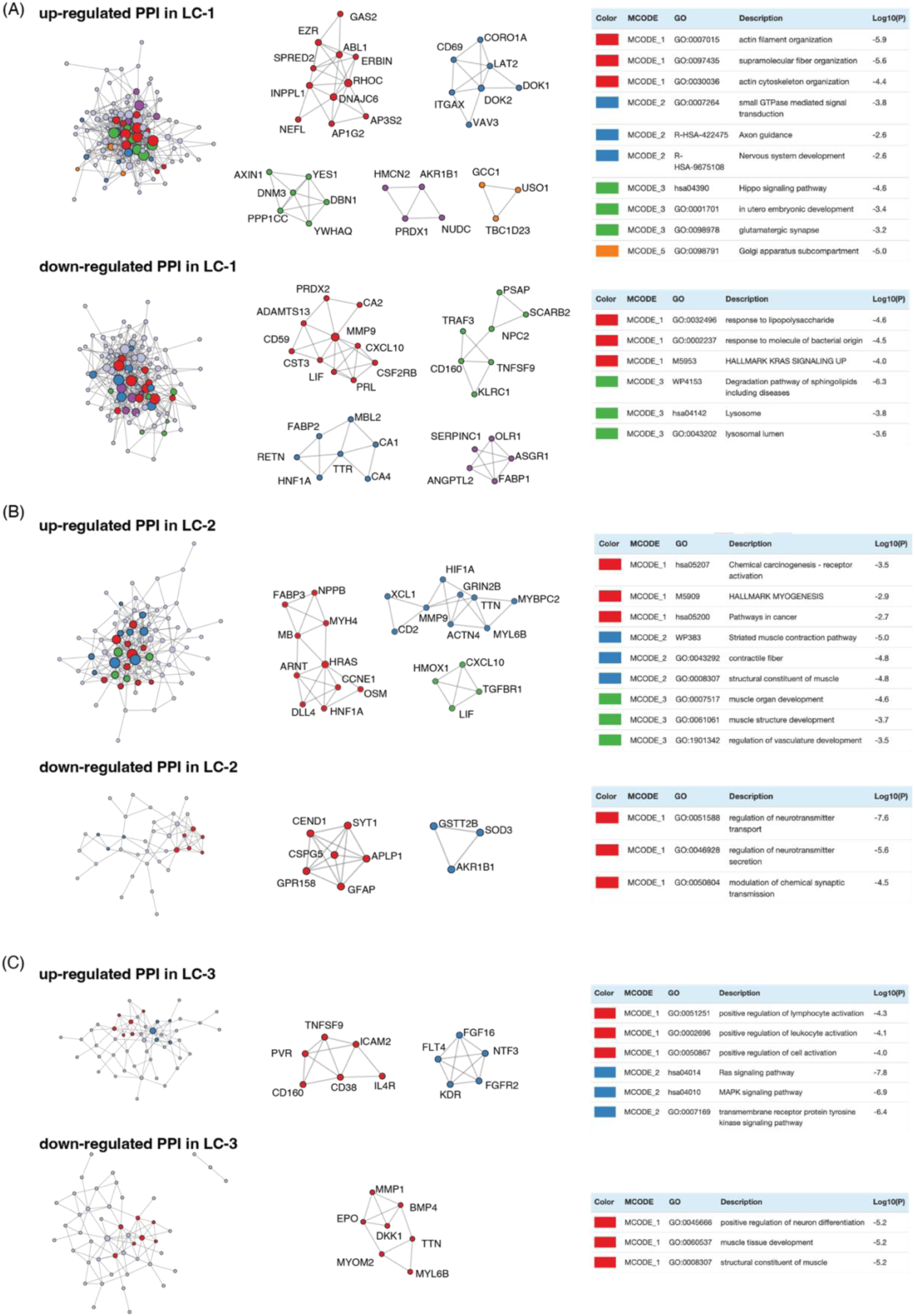
Differential analysis of plasma proteins. Protein-protein interaction (PPI) enrichment analysis for the differentially regulated proteins revealed elevated and suppressed protein-protein interactions in three Long COVID (LC) patient subgroups **(A)** LC-1, **(B)** LC-2, and **(C)** LC-3.

### Human IgG intraperitoneal injection leads to tissue human IgG accumulation in mice

We hypothesized that if autoantibodies are causative in Long COVID IgG antibodies, part are likely to be xenogenetically cross-reactive resulting in tissue damage and subsequent associated symptoms in mice. We purified IgG from patients and pooled them according to the Long COVID subgrouping, obtaining three different human IgG pools. We also included an age-and-sex matched healthy control group sampled prior to the SARS-CoV-2 pandemic (Supp. Table 4). Pooled purified human IgG (hIgG) from each of the three Long COVID subgroups and the HC group were injected into 8 mice per group (4 females and 4 males) at a single dose of 260 mg/kg, which is approximately 1/3 of total circulating IgG per mouse [26]. At day 15 post injection, we assessed the presence of hIgG in these mice across various tissues. We examined hearts, skeletal muscles, spinal cords, and dorsal root ganglia (DRG), as these tissues have been postulated to be affected in Long COVID pathology. Additionally, in spinal cords, we targeted the dorsal horns, where sensory neurons’ afferent nerves synapse with interneurons to process sensory information, including pain perception, a prevalent symptom in Long COVID. We detected hIgG in all examined tissue types, including muscular tissues (heart and skeletal muscles, Fig.S3), spinal cords (Fig.S4), and DRGs (Fig.S5). The intensity of hIgG in mice injected with hIgG from healthy donors (M-HC) was not significantly different from those injected with hIgG from Long COVID patients (M-LC) (Fig. S3-5). We observed some co-localization of hIgG with neurons and glial marker GFAP. No clear differences were found in spinal cords and DRGs between M-HC and M-LC mice (Fig.S6-7). While elevated GFAP expression represents astroglial activation during neurodegeneration [38], we did not observe changes in GFAP expression or co-localization in spinal cords of M-LC compared to M-HC (Fig.S6). Together, these data indicate that the injected hIgG, either from LC or HC groups, were detectable in all the tissues tested at similar levels.

### Long COVID IgG transfer induces sensory hypersensitivity in mice

Next, since neurosensory symptoms, particularly pain, are prevalent in Long COVID patients [3], we investigated whether the detected human IgG in nervous system causes neurological symptoms. To evaluate pain-associated behavior, we measured mechanical sensory thresholds with the von Frey test and thermal sensitivity using the Hargreaves test [27]. Mice that received Long COVID patient IgG developed a pronounced reduction in mechanical sensory threshold (increased mechanical sensitivity) that lasted for at least 15 days compared to the M-HC (Fig. 5A). Considering the Long COVID subgroups, the reduction in mechanical threshold primarily occurred in M-LC1 and M-LC3 (Fig. 5B). Notably, the time course for the development of mechanical hypersensitivity was different in M-LC1 and M-LC3: M-LC1 developed mechanical hypersensitivity from day 3, whilst the mechanical threshold in M-LC3 were already reduced at 24 hours post-hIgG injection.

**Figure 5.**
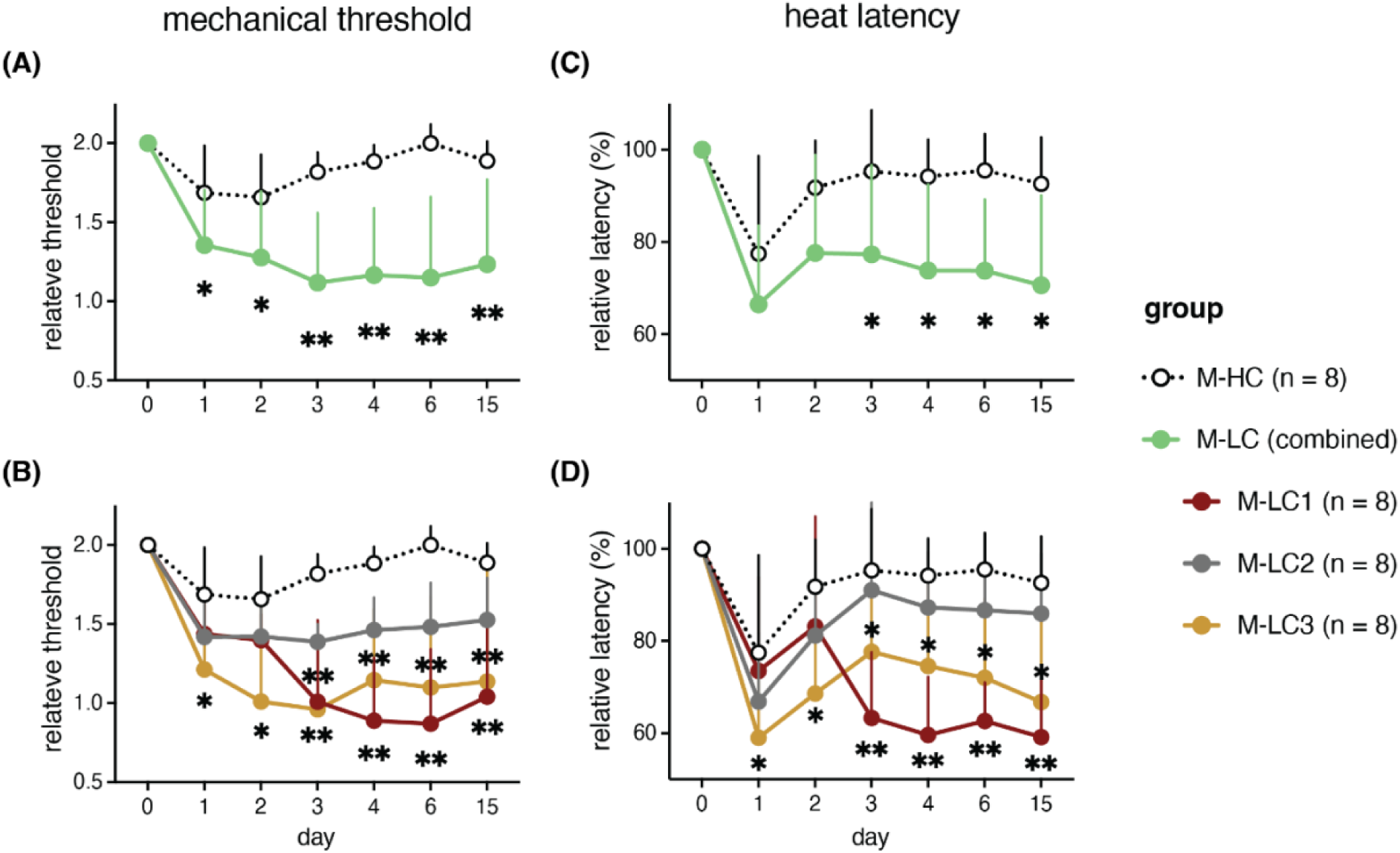
IgG from Long COVID patients induces sensory hypersensitivity in mice. **(A-B)** relative mechanical sensitivity (50% threshold of the mechanical force) to baseline (pre-injection) in the von Frey test. **(C-D)** relative heat sensitivity (latency) to baseline (pre-injection) in the Hargreaves assay. Data points are shown as the mean ± SD. Statistical significance is calculated with a linear mixed effects model with post-hoc comparison using the emmeans package with BH adjustment. *p<0.05, **p<0.01.

In both M-LC and M-HC, the latency to heat stimulation was reduced, which indicates increased heat sensitivity. While this thermal hypersensitivity normalized within 2-3 days in mice injected with HC hIgG, in mice that had received Long COVID hIgG, the hyper-sensitive state persisted until at least 15 days post-injection (the last measurement, Fig. 5C). Considering Long COVID subgroups, the latency to heat stimulation was only persistently reduced in M-LC1 and M-LC3, whereas M-LC2 mice did not significantly differ from the M-HC group at any measured time points (Fig. 5D). Interestingly, mice injected with hIgG from LC-1 group only differed from HC starting from day 3, whilst mice injected with IgG from LC-3 developed thermal hypersensitivity that was stronger than in mice injected with HC igG starting from day 1 after injection. These tests indicate that IgG from different subgroups of Long COVID patients elicited distinct sensory symptoms, with subgroup-specific course in pain-associated behavior.

### Long COVID IgG transfer affects locomotor activity in mice

Considering the wide range of musculoskeletal complications observed in Long COVID patients [7, 39], we hypothesized that transferring Long COVID IgG could also result in impaired movement behavior. We assessed general locomotor activity levels using an open field test [30]. We observed a minor reduction in locomotor activities in mice injected with IgG from Long COVID patients (Fig. 6A). Considering the subgroups, interestingly, only M-LC2 showed significantly reduced walking distance, approximately 40% less than M-HC at one day post-injection (Fig. 6B). In contrast, M-LC1 and M-LC3 did not differ from M-HC at any of the tested time points. Analysis of the moving patterns, indicated that the reduced motor activity in M-LC2 was likely due to an increase in immobility within this group (Fig. 6C-D). In addition to locomotor activity, motor strength, coordination and balance are also important factors in movement behavior. We assessed these motor functions in mice using two rotarod paradigms [40]: one with fixed speed and another with accelerating speed over time to investigate baseline motor coordination and detect subtle impairments in motor coordination, respectively. In both assays, M-LC or M-HC mice did not perform differently in these tests (Fig. 7A-D). Together, this data indicates that transfer of IgG of LC-2 to mice induced transient alterations in general locomotor activity due to increased immobility, without affecting gross coordination and balance.

**Figure 6.**
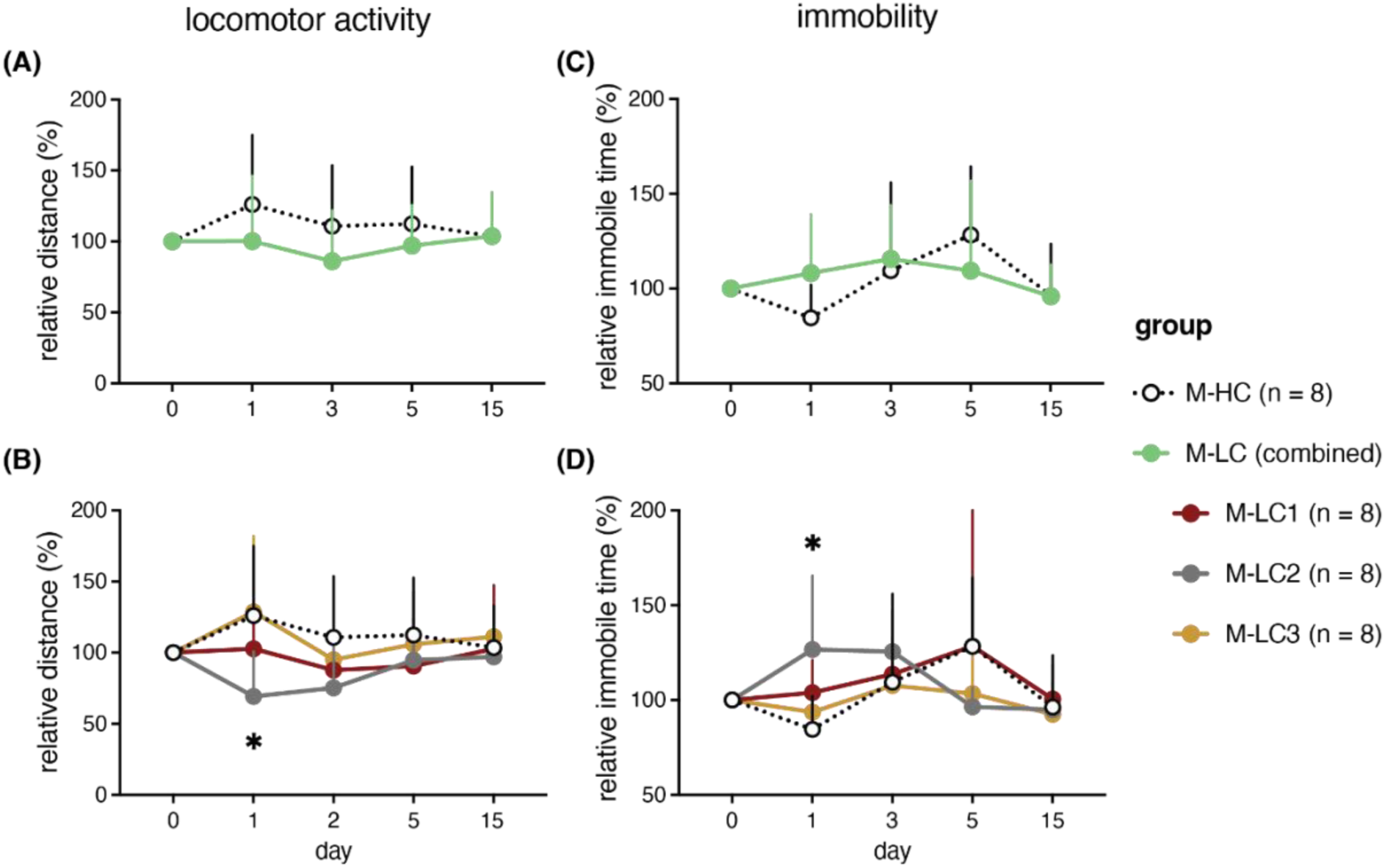
IgGs from Long COVID patient subgroups reduce locomotor activity in mice. Open field test of mice injected with IgG for healthy controls (M-HC) of Long COVID patients (M-LC) **(A-B)** relative walking distance to baseline (pre-injection) in the open field test **(C-D)** relative immobility (time) to baseline (pre-injection) in the open field test. Data points are shown as the mean + SD. Statistical significance is calculated with a linear mixed effects model with post-hoc comparison using the emmeans package with BH adjustment. *p<0.05

**Figure 7.**
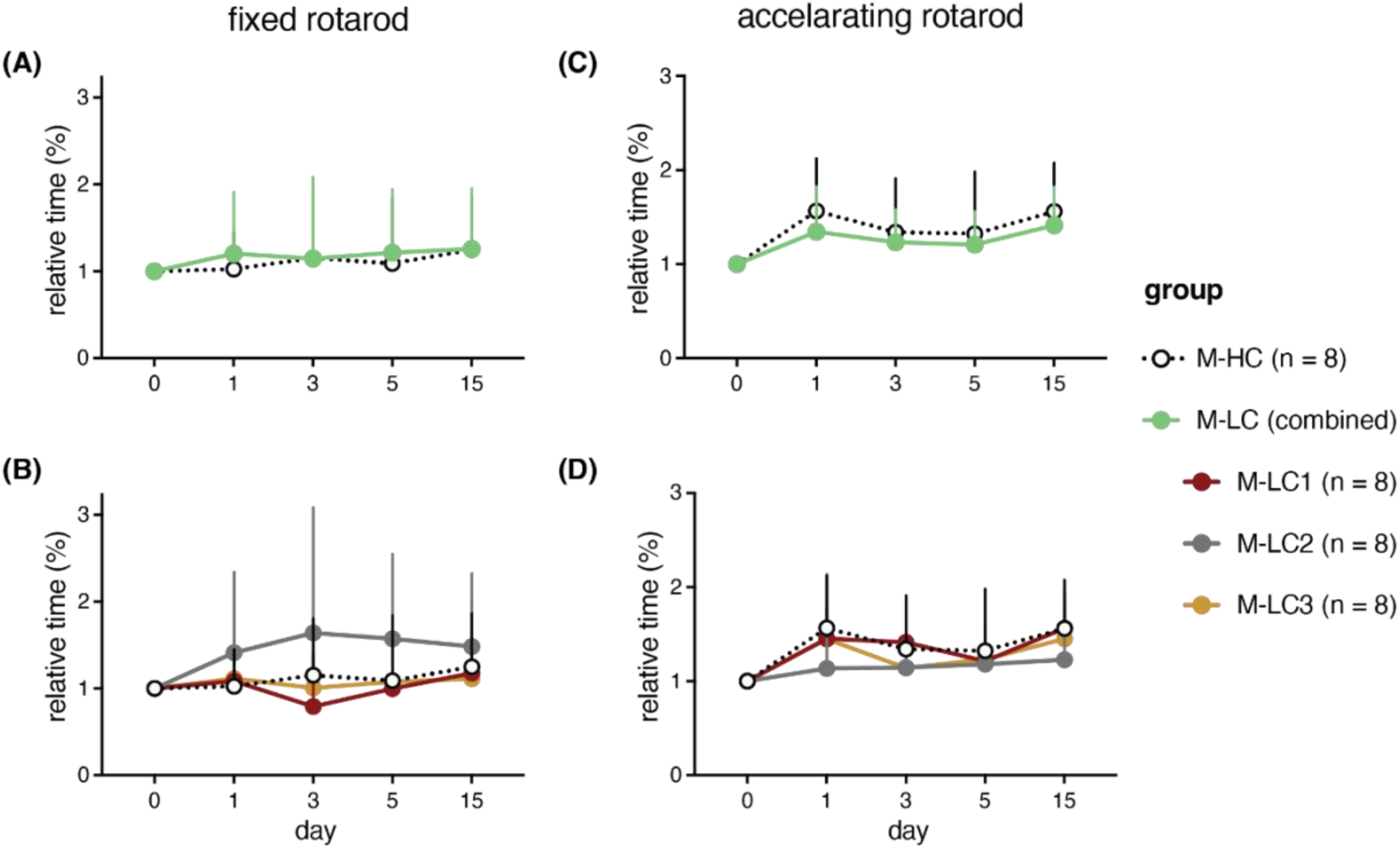
IgG from Long COVID patients did not change time spent on the rotarod. Rotarod test of mice injected with IgG for healthy controls (M-HC) of Long COVID patients (M-LC) **(A-B)** relative time spent on the rotarod in the fixed-speed test **(C-D)** relative time spent on the rotarod in the accelerating test. Data points are shown as the mean + SD. Statistical significance is calculated with a linear mixed effects model with post-hoc comparison using the emmeans package with BH adjustment.

## Discussion

This study investigated the role of IgG in Long COVID pathogenesis. Our findings demonstrate we can identify three distinct Long COVID subgroups using quantitative plasma biomarker analysis. Transfer of IgG from these identified subgroups to mice caused subgroup specific long-lasting pain-associated behavior and transient reduction in locomotor activity, underscoring a causative and heterogeneous role of IgG in Long COVID pathogenesis.

We identified altered plasma proteins in Long COVID patients compared to post-COVID non-Long COVID individuals, including decreased IFN-γ, increased GFAP, and a trend of increased IFN-β. Hypo-IFN-γ [41] and hyper-IFN-α/β [42] during the acute phase has been linked to COVID-19 severity and decreased serotonin levels in later stages, which are associated with Long COVID symptoms [8]. Interestingly, despite mild symptomology during the acute phase, our patients exhibited dysregulated IFN signatures at the time of sampling (over six months post-infection). Further studies are warranted to investigate whether prolonged dysregulation of type-I/II IFNs contributes to Long COVID pathogenesis in patients who did not develop severe COVID-19.

Contrary to previous findings, we did not observe significant differences in acute-phase COVID-19 pro-inflammatory cytokines between non-Long COVID controls and our patients. It is crucial to note that in studies that identified chronic pro-inflammatory states in Long COVID patients, a substantial proportion of the studied population experienced severe illness during the acute phase [32, 43, 44]. In contrast, our cohort exclusively consisted of patients who only had mild symptoms during the acute phase of infection. This distinction underscores the need for further investigation to determine whether Long COVID patients with differing acute-phase severity share the same primary pathophysiology.

Based on GFAP and type-I IFNs, we categorized Long COVID patients into three subgroups, each showing proteome signatures enriched in distinct pathways. These findings suggest various underlying mechanisms trigger different local/systemic tissue damages, leading to the release of corresponding biomarkers into circulation. However, it is possible that these markers are not direct pathogenic factors driving disease progression but rather consequences of Long COVID development and as such may differ based on pathophysiology/symptoms.

Production of autoantibodies is well documented in many viral infections [45], including SARS-CoV-2 [46], influenza [47], West-Nile Virus [48] and Epstein-Barr virus [49]. These autoantibodies contribute to disease severity through various mechanisms, such as inducing cell lysis and activation of leukocytes, complement system, and platelets. Additionally, autoantibodies can act as agonists or antagonists against certain receptors, cytokines, and hormones, thus modulating downstream signaling and neural transmission [50]. Such autoantibody-induced pathology has been extensively described in rheumatic and autoimmune diseases. Recent studies have demonstrated an association between fibromyalgia, ME/CFS, and pathogenic antibodies targeting glia cells [51]. These antibodies have a catalytic function that breaks down myelin basic protein [52], or act as a β2-adrenergic receptor agonist in ME/CFS. Here, we demonstrated that the transfer of IgG from Long COVID patients induced behavioral symptoms in mice, indicating the causal role of IgG in Long COVID pathophysiology. Importantly, the pattern and nature of behavioral changes was different depending on which subgroup of LOC patients the IgG were derived. Mice injected with IgG from LC-1, the patients with elevated plasma levels of neurodegenerative proteins (GFAP and NFL), exhibited the significant increase in mechanical and thermal sensitivity from day 3 to 15 post-intraperitoneal IgG injection. This aligns with the association between astrogliosis, nerve damage and chronic pain [53]. Additionally, MMP1, the top elevated plasma marker in LC-1 patients (Supp. Fig. 3A), can promote sustained pain by inducing neurostructural changes and PAR1 signaling [54, 55]. Mice that received IgG from LC-3 developed a more rapid onset of hypersensitivity post-injection. LC-3 patients’ plasma had higher levels of leukocyte activation proteins, with one of the top upregulated proteins being EIF5A (Supp. Fig. 3C). Intriguingly, inhibition of EIF5A reduces firing in human induced pluripotent stem cells (iPSC)-derived neurons and prevents mechanical hypersensitivity in a model of hyperalgesic priming [56]. M-LC2 mice did not develop significant hypersensitivity to mechanical and heat stimulation. Interestingly, compared to LC-1 and LC-3 patients, the corresponding patients (LC-2) exhibited lower plasma levels of neural proteins, particularly involved in the regulation of neurotransmitter transport and secretion (Fig. 3B). In contrast, LC-2 patients’ plasma was enriched in proteins in skeletal and cardiac muscle pathways. For instance, TTN, one of the top enriched protein in these patients (Fig. 3B), is responsible for the passive elasticity of muscle, and its mutation has been associated with muscle disorders and cardiomyopathies [57]. Mice receiving LC-2 IgG exhibited a significant reduction in distance traveled but without clear motor coordination defects and or grip strength. As motor coordination defects are often associated with neurological pathology, the plasma protein signatures are in line with the observed motor changes in mice that received IgG from LC-2. Moreover, we stratified LC-2 based on increased IFN-β, and high levels of type-I IFN decrease healthy muscle stem cell proliferation [58] and induces contractile weakness, particularly by IFN-β [59]. The potential pathological association between plasma proteome signatures and IgG-induced murine behavioral outcomes underscores the pivotal role of IgG in Long COVID’s heterogeneous pathogenesis. Further investigation is warranted to elucidate if IgGs from individual patients induce varied pathological effects.

Both IgG from Long COVID patients and healthy controls accumulated in satellite cells in the DRG. Notably, a previously published study demonstrated that injecting fibromyalgia patients’ IgG in mice resulted in hIgG accumulation in satellite cells in the DRGs, but did not occur in mice injected with healthy controls’ IgG [24]. It is important to acknowledge that murine Fc receptors may not effectively recognize human antibodies, and human autoantibodies may not recognize the mouse ortholog antigen. These factors can potentially lead to the absence of certain symptomatic manifestations in our mouse models. Additionally, behavioral differences observed in mice may be induced by pathogenic IgGs from specific Long COVID individuals, highlighting the disease’s heterogeneity. Future studies should identify specific pathogenic IgGs, which in turn could contribute to a more nuanced understanding of the disease pathology and possible targeted therapeutic strategies.

In conclusion, our study has demonstrated that passive transfer of IgG from Long COVID patients to mice results in distinct patterns of altered sensory sensitivity and locomotor activity, suggesting the fundamental role of IgG in the heterogenous pathogenesis of Long COVID. Thus, establishing a murine model that mimics Long COVID pathology can present a promising avenue for advancing our understanding of Long COVID pathophysiology and promoting development of targeted and personalized therapeutic interventions.

## Supplementary tables/figures

**Supplementary Figure 1:**
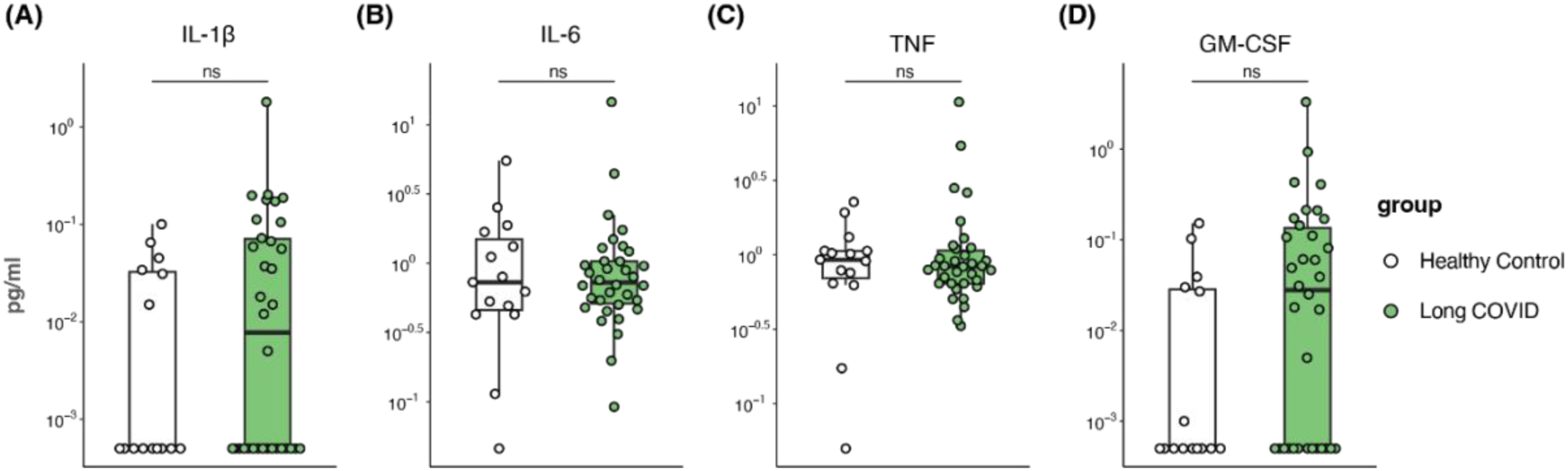
Plasma biomarkers of Long COVID compared to healthy SARS-CoV-2 recovered participants. Targeted quantitative measurements of acute COVID-19 severity markers (IL-1β, Il-6, TNF, and GM-CSF) of 34 Long COVID patients and 15 healthy controls with prior SARS-CoV-2 infection using MSD. All values are shown in pg/ml on a logarithmic y-axis. ns, non-significant.

**Supplementary Figure 2:**
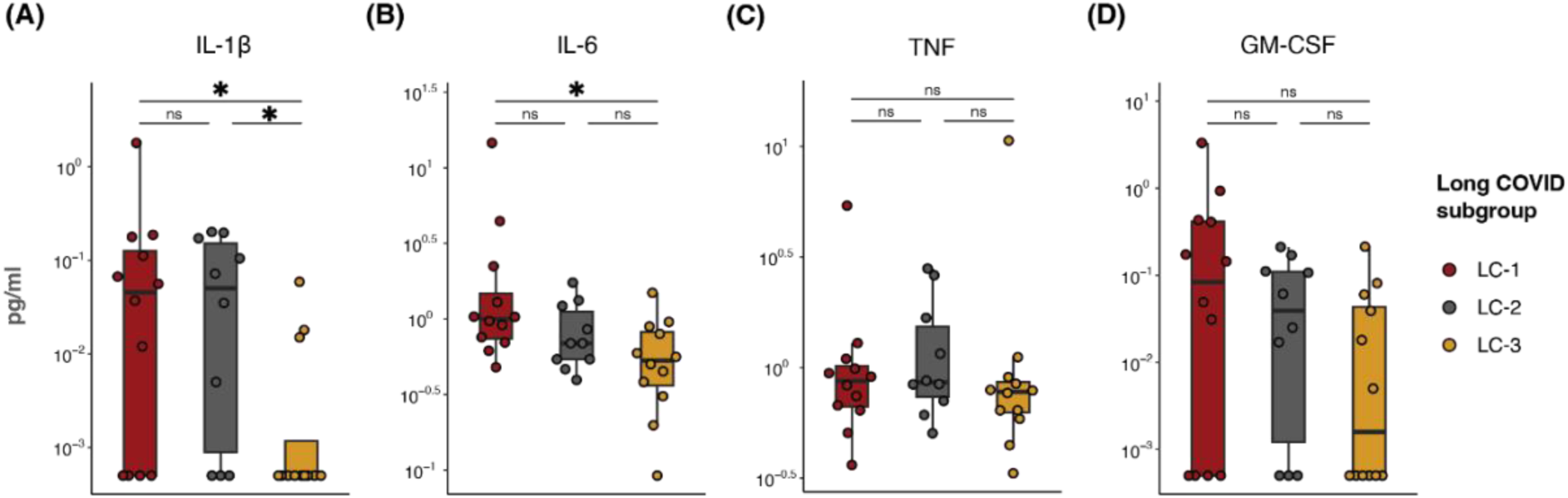
Plasma biomarkers in the Long COVID subgroups. Targeted quantitative measurements of acute COVID-19-associated pro-inflammatory cytokines from 3 Long COVID (LC) patient subgroups (n = 12 for LC1, 10 for LC-2, 12 for LC-3). All values are shown in pg/ml on a logarithmic y-axis. *p<0.05; ns, non-significant.

**Supplementary Figure 3:**
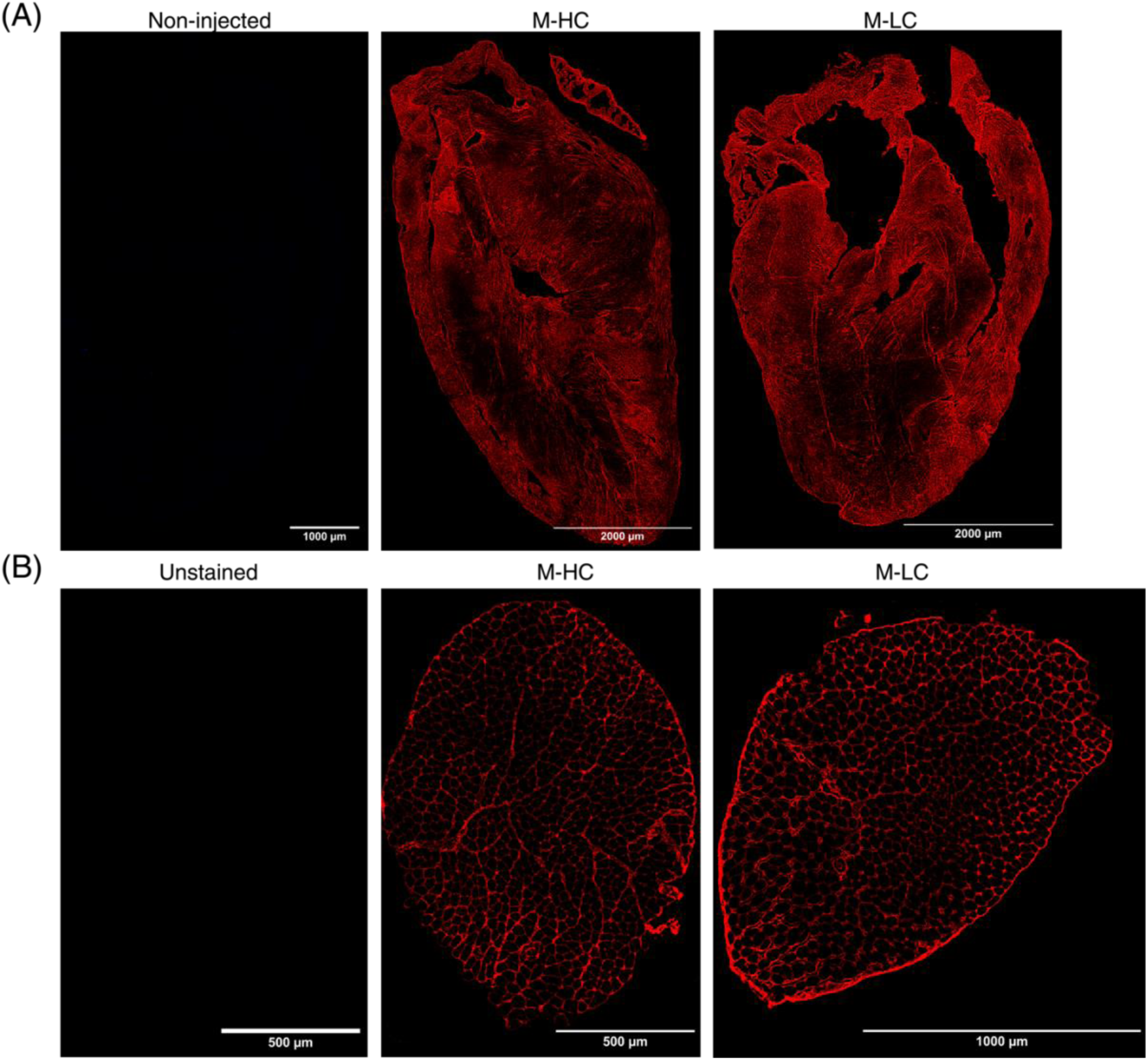
Injected human IgG (hIgG) of Long COVID (LC) patients and healthy donors (HC) localizes in murine heart and skeletal muscles. (A) Representative staining of hIgG (red) in the heart of M-HC (middle panel) and M-LC (right panel). As a negative control, hearts of non-injected mice were included and stained for the presence of hIgG (left panel). (B) Representative staining of hIgG in skeletal muscle of M-HC (middle panel) and M-LC (right panel). As a negative control, unstained skeletal muscle was taken along (left panel).

**Supplementary Figure 4:**
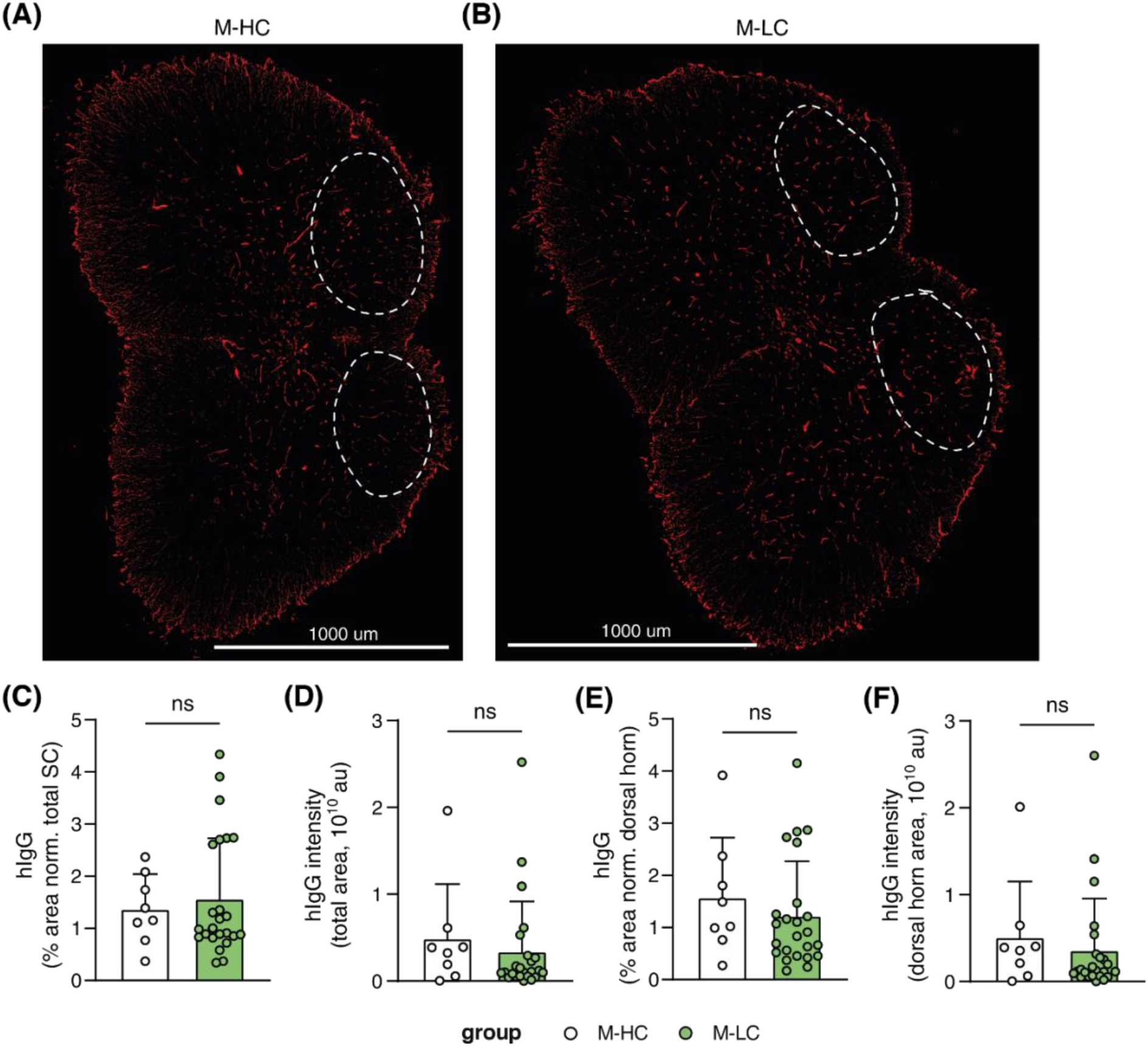
Injected human IgG (hIgG) of Long COVID (LC) patients and healthy donors (HC) detected in murine spinal cord. Representative staining of hIgG (red) from the spinal cord of (A) mice injected with HC IgG antibodies (M-HC) and (B) M-LC. The white dotted line indicates the dorsal horn area. (C) Quantification of hIgG staining area normalized to total spinal cord area per mouse. (D) Quantification of the intensity of hIgG staining in total spinal cord per mouse. (E) Quantification of hIgG staining area, normalized to dorsal horn area per mouse. (F) Quantification of the intensity of hIgG staining in the dorsal horn area. Continuous parametric data were analyzed using t-tests. ns; non-significant.

**Supplementary Figure 5:**
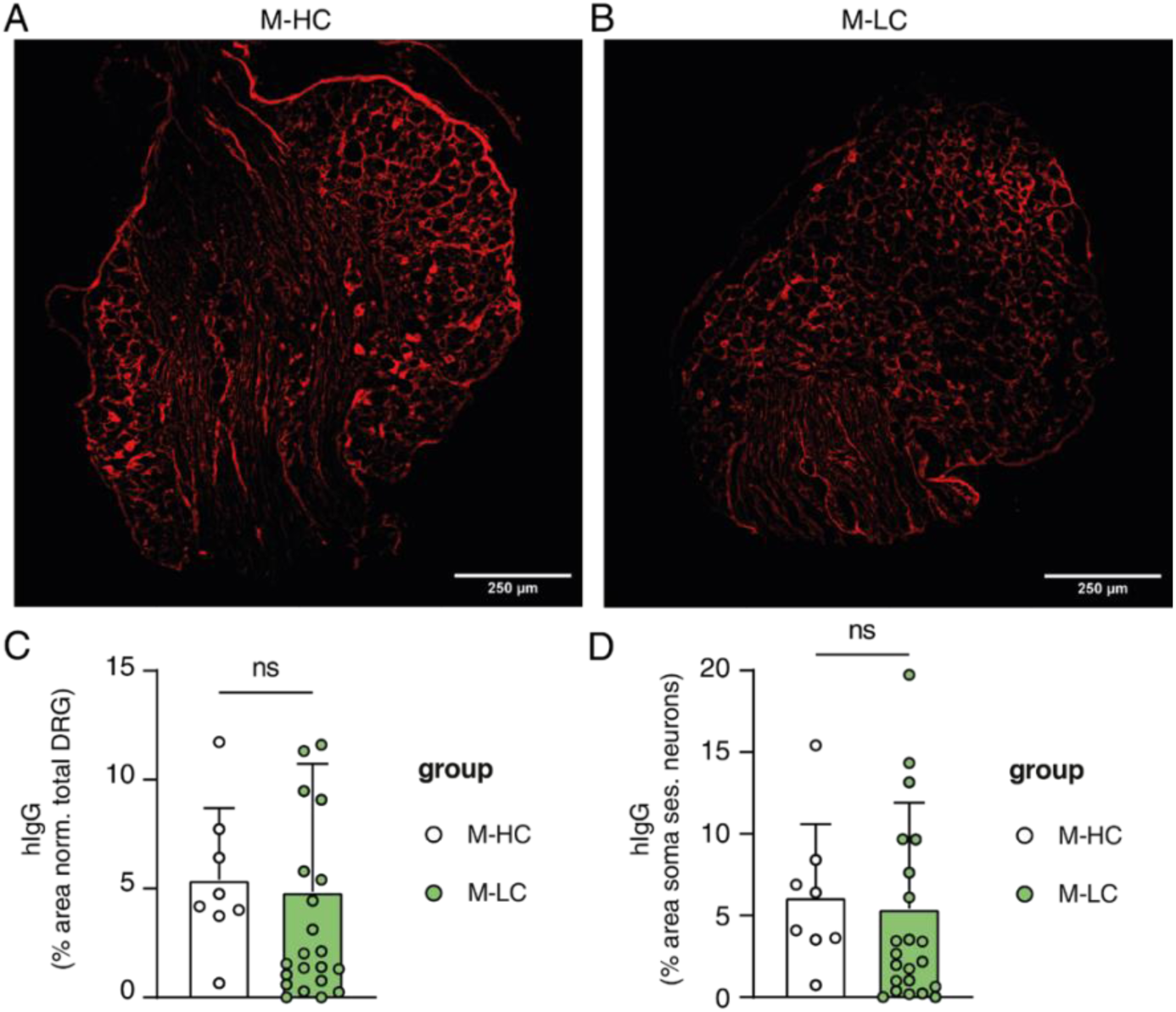
Injected human IgG (hIgG) of Long COVID (LC) patients and healthy donors (HC) detected in murine dorsal root ganglia (DRG). Representative staining of hIgG (red) in the DRG of (A) mice injected with HC IgG antibodies (M-HC) and (B) M-LC. (C) Quantification of the percentage of area in the DRG containing hIgG normalized to total DRG area per mouse. (D) Quantification of the percentage of DRG area that contained hIgG, normalized to the area that contained the somas of sensory neurons per mouse. Continuous parametric data were analyzed using t-tests. ns; non-significant.

**Supplementary Figure 6:**
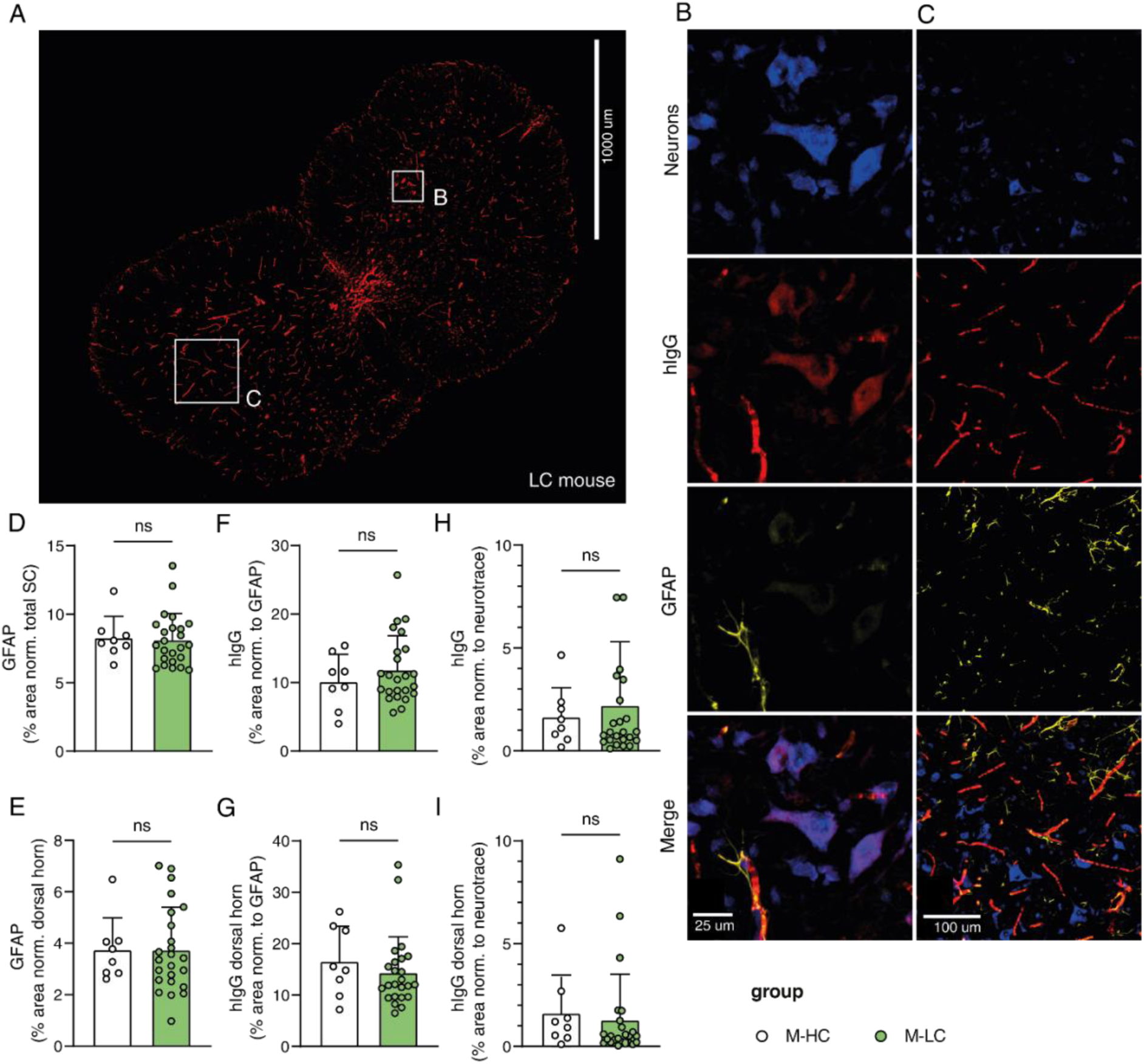
Human IgG (hIgG) of Long COVID (LC) patients and healthy donors (HC) injected into mice, co-localizes with spinal cord neurons and astrocytes. Representative pictures of spinal cords stained for neurons (NeuroTrace, blue), hIgG (red) and astrocytes (glial fibrillary acidic protein (GFAP), yellow). Squares represent regions used to illustrate co-localization in B-C. (B) Magnification of figure A demonstrating hIgG staining which co-localizes with neurons. (C) Magnification of figure A demonstrating hIgG staining which partly co-localizes with astrocytes. (D-E) Quantification of the percentage of the area positive for GFAP for (D) total spinal cord or for the (E) dorsal horn for HC and LC injected hIgG. (F) Quantification of hIgG area that co-localized with GFAP staining in the total spinal cord. (G) Quantification of hIgG co-localization with GFAP staining, normalized to total GFAP+ area in the dorsal horn area. (H) Quantification of hIgG co-localization with NeuroTrace staining, normalized to overall NeuroTrace+ area in the total spinal cord. (I) Quantification of hIgG co-localization with NeuroTrace staining, normalized to total NeuroTrace in the dorsal horn area. Continuous parametric data were analyzed using a t-test. ns; non-significant.

**Supplementary Figure 7:**
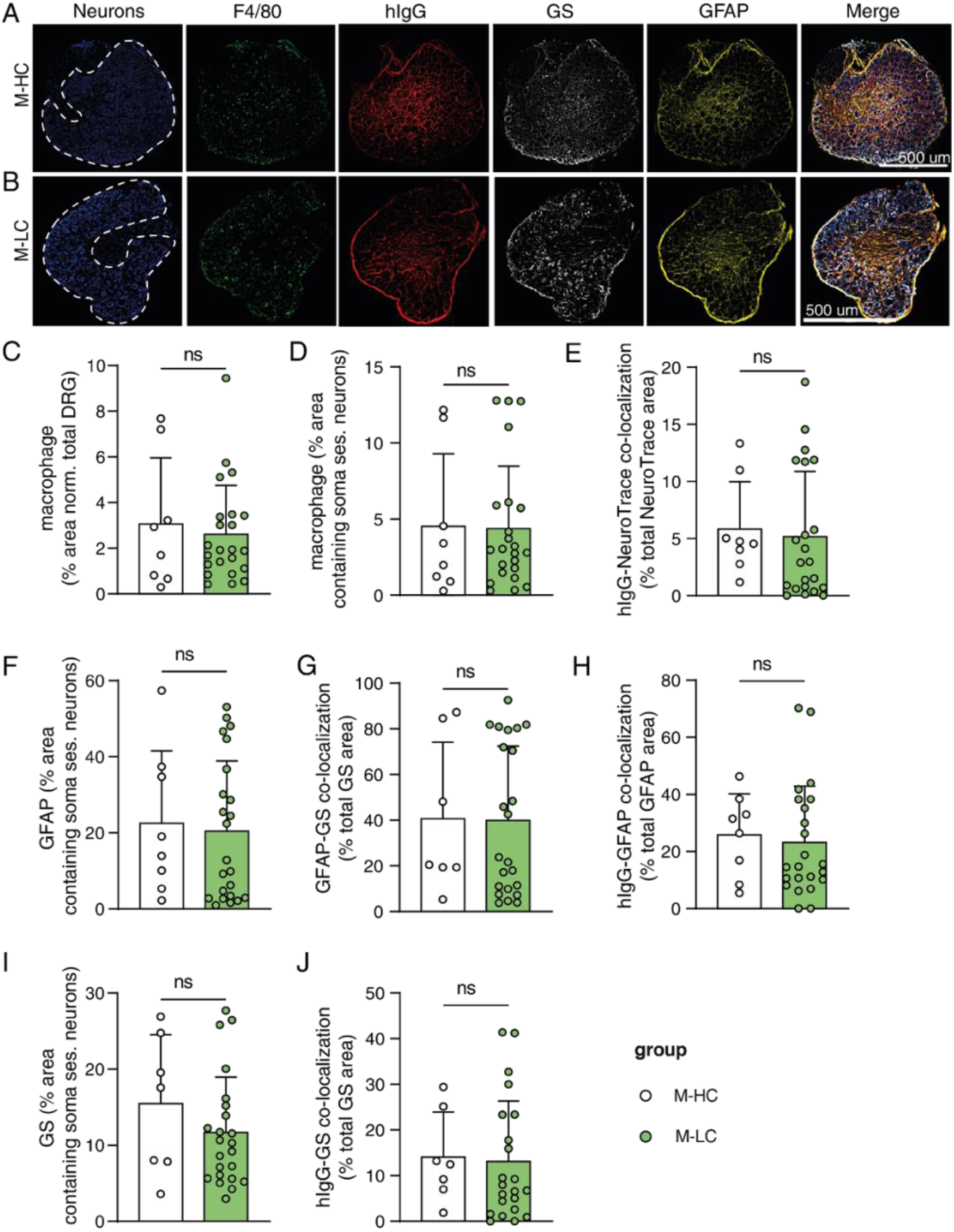
Injected human IgG (hIgG) of Long COVID patients (LC) and healthy donors (HC) detected in murine dorsal root ganglia. (A-B) Representative pictures of dorsal root ganglia (DRG) stained for neurons (NeuroTrace, blue), macrophages (F4/80, green) hIgG (red), satellite cells (glutamine synthetase (GS), white) and Glial Fibrillary Acidic Protein (GFAP, yellow). The white dotted line indicates the area of the DRG containing the somas of sensory neurons, based on NeuroTrace labelling (left panel). Quantification of F4/80 expressing macrophages in (C) total DRG tissue and (D) the soma of sensory neurons of mice injected with HC (white, M-HC) and LC IgG (green, M-LC) antibodies. (E) Quantification of hIgG co-localization with NeuroTrace staining. Quantification of GFAP positive area withing the DRG area containing (F) the somas and (G) GS-positive area. (H) Quantification of hIgG co-localization with GFAP staining, normalized to overall GFAP area in DRG. (I) Quantification of GS in the soma of sensory neurons. (J) Quantification of hIgG co-localization with GS staining, normalized to overall GS area in DRG. Continuous parametric data were analyzed using t-tests. ns; non-significant.

**Supplementary Table 1.** Demography of non-Long COVID post-COVID-19 controls.

|  | Healthy controls<br>n=15 |
| --- | --- |
| Sex = Male (%) | 3 (20.0) |
| Age (median [IQR]) | 35 [32, 49] |
| Charlson comorbidity index (median [IQR]) | - |
| Time from infection to sampling, days (median [IQR]) | 278 [271, 288] |
| Vaccination before initial SARS-CoV-2 infection = Yes (%) | 0 (0) |
| Vaccination before sampling = Yes (%) | 0 (0) |
| Working hours prior SARS-CoV-2 infection (median [IQR]) | - |

**Supplementary Table 2.** Baseline characteristics of long COVID subgroups.

|  | Long COVID-1<br>n=12 | Long COVID-2<br>n=10 | Long COVID-3<br>n=12 | p-value |
| --- | --- | --- | --- | --- |
| Sex = Male (%) | 3 (25.0) | 1 (10) | 2 (16.7) | 0.65 |
| Age (median [IQR]) | 51 [45, 58] | 40 [31, 45] | 42 [34, 46] | 0.08 |
| Charlson Comorbidity Index<br>(median [IQR]) | 1 [0, 2] | 0 [0, 0] | 0 [0, 0] | **0.01** |
| Time from infection to sampling,<br>days (median [IQR]) | 232 [160, 316] | 330 [274, 375] | 231 [190, 381] | 0.15 |
| Vaccination = Yes (%) | 10 (83) | 9 (90) | 11 (91.7) | 0.80 |
| Working hours prior SARS-CoV-2<br>(median [IQR]) | 34 [28, 38] | 32 [32, 43] | 35 [24, 40] | 0.89 |

**Supplementary Table 3.** DEG tables OLINK. See attached supplementary files.

**Supplementary Table 4.** Demography of pre-pandemic controls.

|  | Pooled healthy<br>controls<br>n=34 |
| --- | --- |
| Sex = Male (%) | 6 (17.6) |
| Age (median [IQR]) | 43 [34, 50] |

## Funding

This work was supported by the Patient-Led Research Collaborative of Long COVID (grant ID: C1), Stichting Long-COVID consortium grant, the Netherlands Organization for Health Research and Development (ZonMw, grant number 10430142210001).

The funders had no role in study design, data collection, and analysis, decision to publish, or preparation of the manuscript.

## Conflicts of interest

The authors report no conflict of interest.

## Data sharing

Data can be shared upon reasonable request after approval of a proposal with a signed data access agreement and always in collaboration with the study group.

## Author contribution

Conceptualization: JdD, NE;

Cohort: BA, MvV, MKB, AHAL, AUMC COVID-19 biobank; Methodology: HJC, BA, HLDMW, JP, ML, AB;

Investigation: HJC, BA, HLDMW, JP, CEG, AB, ES, SV, PSSR;

Formal analysis: HJC, BA, HLDMW, JP, AB;

Original draft: BA, HJC;

Review and editing: JdD, NE,WJW, HJC, BA, HLDMW, PSSR, JP, AB, GV;

Funding: BA, JdD, NE

## Acknowledgments

We would like to thank all the patients and the staff of the Amsterdam UMC post-covid clinic for their contribution.

## Group authorships

Amsterdam UMC COVID-19 biobank

Michiel van Agtmael2, Anne Geke Algera1, Brent Appelman2, Floor van Baarle1, Martijn Beudel4, Harm Jan Bogaard5, Marije Bomers2 Peter Bonta5, Lieuwe Bos1, Michela Botta1, Justin de Brabander2, Godelieve de Bree2, Sanne de Bruin1, Marianna Bugiani5, Esther Bulle1, David T.P. Buis1, Osoul Chouchane2 Alex Cloherty3, Mirjam Dijkstra12, Dave A. Dongelmans1, Romein W.G. Dujardin1, Paul Elbers1, Lucas Fleuren1, Suzanne Geerlings2 Theo Geijtenbeek3, Armand Girbes1, Bram Goorhuis2, Martin P. Grobusch2, Laura Hagens1, Jorg Hamann7, Vanessa Harris2, Robert Hemke8, Sabine M. Hermans2 Leo Heunks1, Markus Hollmann6, Janneke Horn1, Joppe W. Hovius2, Katja de Jong2, Menno D. de Jong9, Rutger Koning4, Bregje Lemkes2, Endry H.T. Lim1, Niels van Mourik1, Jeaninne Nellen2, Esther J. Nossent5, Sabine Olie4, Frederique Paulus1, Edgar Peters2, Dan A.I. Pina-Fuentes4, Tom van der Poll2, Bennedikt Preckel6, Jan M. Prins2, Jorinde Raasveld1, Tom Reijnders2, Maurits C.F.J. de Rotte12, Michiel Schinkel2, Marcus J. Schultz1, Femke A.P. Schrauwen12, Alex Schuurman2, Jaap Schuurmans1, Kim Sigaloff1, Marleen A. Slim1,2, Patrick Smeele5, Marry Smit1, Cornelis S. Stijnis2, Willemke Stilma1, Charlotte Teunissen11, Patrick Thoral1, Anissa M Tsonas1, Pieter R. Tuinman1, Marc van der Valk2, Denise Veelo6, Carolien Volleman1, Heder de Vries1, Lonneke A. Vught1,2, Michèle van Vugt2, Dorien Wouters12, A. H (Koos) Zwinderman13, Matthijs C. Brouwer4, W. Joost Wiersinga2, Alexander P.J. Vlaar1, Diederik van de Beek4.

1. Department of Intensive Care, Amsterdam UMC, Amsterdam, The Netherlands;

2. Department of Infectious Diseases, Amsterdam UMC, Amsterdam, The Netherlands;

3. Experimental Immunology, Amsterdam UMC, Amsterdam, The Netherlands;

4. Department of Neurology, Amsterdam UMC, Amsterdam Neuroscience, Amsterdam, The Netherlands;

5. Department of Pulmonology, Amsterdam UMC, Amsterdam, The Netherlands;

6. Department of Anesthesiology, Amsterdam UMC, Amsterdam, The Netherlands;

7. Amsterdam UMC Biobank Core Facility, Amsterdam UMC, Amsterdam, The Netherlands;

8. Department of Radiology, Amsterdam UMC, Amsterdam, The Netherlands;

9. Department of Medical Microbiology, Amsterdam UMC, Amsterdam, The Netherlands;

10. Department of Internal Medicine, Amsterdam UMC, Amsterdam, The Netherlands;

11. Neurochemical Laboratory, Amsterdam UMC, Amsterdam, The Netherlands;

12. Department of Clinical Chemistry, Amsterdam UMC, Amsterdam, The Netherlands;

13. Department of Clinical Epidemiology, Biostatistics and Bioinformatics, Amsterdam UMC, Amsterdam, The Netherlands.

## Notes

### Competing Interest Statement

The authors have declared no competing interest.

